# UV-induced feather color change reflects its porphyrin content

**DOI:** 10.1101/2023.05.14.540673

**Authors:** Masaru Hasegawa, Emi Arai, Shosuke Ito, Kazumasa Wakamatsu

## Abstract

Pigmentary coloration is widespread in animals. Its evolutionary and ecological features are often attributed to the property of predominant pigments; therefore, most research has focused on predominant pigments such as carotenoids in carotenoid-based coloration. However, coloration results from predominant pigments and many other minority pigments, and the importance of the latter is overlooked. Here, we focused on porphyrin, an “uncommon” pigment found in bird feathers, and investigate its importance in the context of feather color changes in the barn swallow *Hirundo rustica*. We found that the “pheomelanin-based coloration” of the barn swallow faded after the irradiation of UV light, and this effect was particularly strong in the feathers of young swallows (nestlings and fledglings, here). We also found that it is not the predominant pigment, pheomelanin, but protoporphyrin IX pigment that showed the same pattern of depigmentation after the irradiation of UV light, particularly in the feathers of young swallows. In fact, the abovementioned age-dependent feather color change was statistically explained by the amount of porphyrin in the feathers. The current study demonstrates that a minority pigment, porphyrin, explains within-season dynamic color change, an ecological feature of feather coloration. The porphyrin-mediated rapid color change would benefit young birds, in which feather coloration affects the parental food allocation during a few weeks before independence, but not later. Future studies should not ignore these minor but essential pigments and their evolutionary and ecological functions.

**Significance statement:** Predominant pigments are assumed to determine animal coloration and its ecological features. It is then not surprising that the evolutionary and ecological features of animal coloration are often attributed to the chemical properties of predominant pigments. However, coloration results from predominant pigments and several other minority pigments. By irradiating UV light on reddish throat feathers of the barn swallow *Hirundo rustica*, we examined within-season dynamic color change in relation to a minority pigment, porphyrin, which has not previously been examined but is a candidate pigment for feather color change, because porphyrin can be easily photodegraded. We found that not the predominant pigment, pheomelanin, but porphyrin pigments explained the feather color change. Minor pigments and their chemical properties should not be dismissed to understand the ecological and evolutionary functions of animal coloration.

## Introduction

Pigmentary coloration is widely distributed in animals and serves many functions including camouflage, interspecific signaling (e.g., warning coloration), and intraspecific signaling (e.g., mate attraction; reviewed in Andersson 1994; Espmark et al. 2000; Hill and McGraw 2006). Pigmentary coloration is often subdivided by predominant pigments, such as carotenoid-based coloration, eumelanin-based coloration, and pheomelanin-based coloration (e.g., see McGraw 2006a,b on bird coloration). It is then not surprising that the evolutionary and ecological features of animal coloration are often attributed to the chemical properties of predominant pigments. For example, carotenoid-based coloration can serve as an honest signal of quality to signal receivers, because carotenoids consumed as antioxidants cannot simultaneously serve as feather colorants (reviewed in McGraw 2006a; also see McGraw 2005 for other major pigments). While such a simple explanation is appealing, coloration results from predominant pigments and several other minority pigments. The evolutionary and ecological importance of the latter is ignored, solely because they are less abundant than the former.

Porphyrin is a class of endogenously synthesized colorants that give a reddish, rusty, or brown appearance (McGraw 2006c). Many animals manufacture porphyrins, as evident from the fact that a well-known blood pigment, hemoglobin, is synthesized from natural porphyrin via the addition of iron (i.e., iron-protoporphyrin IX; McGraw 2006c). Nevertheless, besides some exceptions (e.g., the European hedgehog *Erinasceus europaeus*), birds are the only animals that can deposit porphyrin in integumentary organs such as skin and feathers (Galván et al. 2016). Although feather porphyrin and its fluorescent nature is well-known in certain non-passerine birds, such as owls and nightjars, passerines also deposit porphyrin into their feathers (e.g., Ozaki and Imamura 2019). The function of feather porphyrin has not been known (reviewed in McGraw 2006c), but it might function as an “ephemeral” signal because porphyrin is easily photodegraded (Galván et al. 2016). By depositing porphyrin rather than more stable alternatives, such as pheomelanin pigments, which also provide a rusty or brown appearance (McGraw 2006b,c), birds might display conspicuous coloration for only a short period and may even directly signal the freshness of the feathers (Galván et al. 2016). However, the effect of porphyrin on feather color change, particularly in its importance relative to predominant feather pigments (e.g., pheomelanin), remains to be clarified.

In the current study, we investigated porphyrin pigmentation in the context of feather color change using the throat feathers of the barn swallow *Hirundo rustica*. Throat feather coloration of the barn swallow is reported to be used in mate choice (e.g., Ninni 2003; Safran et al. 2005), intrasexual contests (e.g., Hasegawa et al. 2014; Wilkins et al. 2015), and parental preferential provisioning of their young (e.g., Romano et al. 2016; Arai et al. unpublished data). Throat feather coloration mainly consists of pheomelanin and eumelanin pigments (e.g., McGraw et al. 2005), and this coloration changes over time (e.g., Hasegawa et al. 2008; Safran et al. 2010; Arai et al. 2015). Moreover, the color change is reported to be associated with the quality of birds (e.g., Safran et al. 2010), and, within-season changes in feather coloration affect the response of signal receivers (e.g., Safran et al. 2005). In this species, feather color change is particularly notable in juveniles, because their initially pinkish feathers fade to whitish before the next molt so that old and newly molted feathers are visually distinguishable (e.g., see Fig. S1 for an example). Previous studies often assume that pheomelanin and eumelanin pigments are the main determinants of feather coloration and its evolutionary and ecological features, including feather color change (e.g., Arai et al. 2015, 2017, 2019). However, this “pheomelanin-based” coloration might be affected by other pigments, such as porphyrin, which has not previously been examined but is a candidate pigment for feather color change, particularly in the young, in which the feather color change seems striking (see above).

Here, by irradiating UV light (peak: 365 nm) on feathers for 30 days, we investigated feather color change and its relationship with porphyrin pigmentation in the barn swallow. We predicted that, if porphyrin, which can be easily photodegraded, contributes to feather color change in this study system, feather coloration would be affected by UV treatment, particularly in the young (see above). The effect of UV light would be particularly noticeable when examining the feather color change before and after the experiment. The amount of porphyrin in the feathers would show a pattern similar to that of feather coloration. Then, this pigment (but not eumelanin and pheomelanin pigments) would explain the variation of the feather color change. Lastly, to clarify whether or not age-difference in porphyrin content in feathers, if any, is caused by the time elapsed after feather emergence, we examined porphyrin content in the feathers collected from molting adults during the non-breeding season. This was then compared with those collected from adults and young during the breeding season. If the amount of porphyrin is simply determined by the time elapsed after feather emergence, the feathers of molting adults would contain a relatively large amount of porphyrin similar to those of young. Based on previous studies (e.g., McGraw 2006c; Galván et al. 2016, 2018), we focused on coproporphyrin III and protoporphyrin IX as candidate porphyrin pigments. We discuss the observed patterns from the evolutionary and ecological perspectives.

## Methods

### Field survey

The field survey was conducted during the breeding season (1 April–3 July) in 2022 in a residential area of Tsurugi Town, Ishikawa Prefecture, Japan (36°26′N, 136°37′E). We inspected nests every third day to record breeding events such as laying dates, clutch size, and hatching dates. Adults were captured in sweep nets while roosting at night. Each bird was provided with a numbered aluminum ring and a unique combination of half-sized colored rings. The sex of each individual was determined by the tail shape and by the presence (female) or absence (male) of an incubation patch (Turner 2006). During the field survey, we also captured nestlings by hand from breeding nests. Upon capture, throat feathers (approximately 20–40 feathers per bird) were plucked from each bird. In the current study, we used throat feathers of adult females, simply because the capture of females was relatively easier (e.g., Safran and McGraw 2004), and to exclude the potential confounding effect of sex difference in pigmentation (e.g., see Arai et al. 2019 for Japanese swallows). All adult females completed their molt. We found that nestling throat feathers were too small to accurately measure coloration when they were young (see Arai et al. 2017 for detailed feather development in Japanese barn swallows), and thus, concerning nestlings, we plucked throat feathers from well-grown, 16–18-day-old nestlings. Nestling sex was not identified, because the sex difference in throat coloration is at best small (Arai et al. 2018). In this population, we collected samples from seven different females and seven different nestlings. The seven nestlings were captured from seven different breeding nests (i.e., samples were collected from one nestling per nest).

To increase sample size, feather samples were also collected from a breeding population in Miyazaki City, Miyazaki Prefecture, Japan (31°54′N, 131°25′E). As in the Tsurugi population, adults were captured in sweep nets in this population. In addition to adult females, we also captured fledglings, instead of nestlings, in sweep nests, because most nestlings fledged before the study period (5–8 July, 2022). In this population, samples were collected from three different females and three different fledglings. The three fledglings were captured at different nests and roosting sites.

Some barn swallows remain year-round in the Miyazaki population (Hasegawa 2020), and thus newly molted feathers can be collected from molting swallows, in which old and newly molted feathers can be easily distinguished (Fig. S1). For this reason, during 12–17 February 2023, wintering swallows were captured there to collect newly molted throat feathers. We collected samples from three different molting adults, although we could not identify the sex of these individuals (because wintering swallows do not develop brood patches, thereby preventing successful sex identification, particularly when their feathers are still growing).

### Color measurements

In the laboratory, we piled five throat feathers collected from each individual on a piece of white paper so that the perimeters of the feathers coincided. From each bird, two feather samples were taken, one for the control and the other for the UV treatment group (see below). The feather samples were scanned at 800 dpi resolution using a scanner (GT X970; Epson, Tokyo, Japan), and the images obtained were imported into Photoshop Elements 15 (Adobe Systems, San Jose, CA). We measured the mean red-green-blue (RGB) values for a 30 × 30 pixel square near the distal end of the feather sample. The mean RGB values were converted into hue-saturation-brightness (HSB) values, using the algorithm described by Foley and van Dam (1984). Throat feathers exhibit few ultraviolet reflectance, so this method captured most of the variation in throat coloration (e.g., Safran and McGraw 2004). Because previous studies in this study system also used HSB values (e.g., McGraw et al. 2005; Hasegawa et al. 2008; Arai et al. 2015), this method enables us to directly compare our results with those of previous studies. Details of the methodology are described elsewhere (Hasegawa et al. 2008).

### Experimental UV irradiation

The experiment was conducted from 27 October to 28 November 2022 with a short pause to check the progress of the experiment. Before the start of the experiment, we measured the coloration of all 40 feather samples (see above). Two feather samples taken from the same birds resembled each other, demonstrating high repeatability of HSB values between samples (Hue: repeatability = 0.92, *F_19,20_* = 25.34, *p* < 0.0001; Saturation: repeatability = 0.96, *F_19,20_* = 55.56, *p* < 0.0001; Brightness: repeatability = 0.93, *F_19,20_* = 26.84, *p* < 0.0001). Thus, between-individual variation was much larger than within-individual variation in our measurements. Two feather samples each were taken from 20 different individuals, one for the control and the other for the UV treatment group (see above), and thus control and UV treatment groups each had 20 feather samples. Regarding the UV treatment group, feather samples were placed in a closed cardboard box (size: 37 cm × 25 cm × 14 cm), in which they were continuously exposed to UV light (FL365-SD, 27W; peak spectrum: 365 nm, approximate range: 350–400 nm, intensity: 1300 µW/cm^2^ from a distance of 15 cm) at room temperature for 30 days. The intensity of the UV light was within the range of natural UV radiation in Japan (e.g., the intensity of UV-A radiation in open fields between 900 and 1600 in August was 1000–2500 µW/cm^2^ in Akashi City, Hyogo Prefecture in Japan: Ishiuchi et al. 2012). Regarding the control group, samples were placed in the same-sized cardboard box without UV light (i.e., they were held in complete darkness) at room temperature for 30 days. The two treatment groups experienced the same amount of time in the boxes, because the two treatments started and ended at the same time. The two boxes were held in the same room. After the end of the experiment, the coloration of all feather samples was measured again, and the pigment content of each feather sample was evaluated using high-performance liquid chromatography (HPLC; see below).

### Pigment quantification via HPLC

We followed the extraction protocols of porphyrins from feathers given in preceding studies (Galván et al. 2016, 2018; Loules et al. 2022). In short, five feathers (0.2–0.3 mg) were extracted in 300 µL 6 M HCOOH plus acetonitrile (1:1), by homogenizing them with a Tenbroeck tissue grinder. Notably, unlike in the previous study (e.g., Arai et al. 2017), we did not cut off the feathers’ basal part to effectively extract protoporphyrin pigments. After centrifugation in a glass conical tube, the supernatants (50 µL) were subjected to the following HPLC analyses.

We used the precipitates from the feather extraction to analyze eumelanin (pyrrole-2,3,5-tricarboxylic acid; PTCA) and pheomelanin (thiazole-2,4,5-tricarboxylic acid; TTCA). Precipitates (and homogenizer) were washed once with 1 mL water by centrifugation, dried in a desiccator, and subjected to alkaline hydrogen peroxide oxidation (AHPO) as described in previous research (Ito et al. 2011, 2020) using 40% volumes of reagents (Arai et al. 2018, 2019).

The following HPLC conditions were applied. First, A Waters (Milford, MA, USA) Spherisorb ODS 2 (5 µm particle size, 4.6 mm × 250 mm) HPLC column was used. Then, protoporphyrin IX (PPIX) and coproporphyrin III (CPIII) (standards: 1 µg/mL 6 M HCOOH plus acetonitrile (1:1)) were analyzed by isocratic elution under different methanol concentrations. PPIX was analyzed by methanol and 1 M ammonium acetate = 90:10 (v/v). We used high concentrations of ammonium acetate to ensure high buffering capacity. The column temperature was 70℃ (we use the high temperature to make the PPIX peak sharp and high). VIS detection was performed with a UV/VIS detector at 402 nm (Galván et al. 2016). PPIX appeared at 7.5 min. CPIII was analyzed separately by methanol and 1M ammonium acetate = 60:40 (v/v) at a column temperature of 50 ℃. CPIII appeared at 6.9 min.

### Statistical procedures

A linear mixed-effects model (lmer function in the R package lme4: Bates et al. 2015) was used to test the effects of treatment (“Control” and “UV treatment” were denoted as 0 and 1, respectively, and “adults” and “young” were denoted as 0 and 1, respectively) and the interaction between treatment and age on each dependent variable (i.e., hue, saturation, and brightness for color measurements, their changes before and after the experiment, and the amount of each pigment). As before (e.g., Arai et al. 2017), the amount of pigmentation was log-transformed before analysis to fit a normal distribution (Shapiro-Wilk test; before: 0.66 < *W* < 0.96; after: *W* > 0.93). We also examined plumage coloration in relation to pigmentation to test the effect of pigmentation on coloration. In these models, bird identity was included as a random factor to control for pseudoreplication. We did not include the identity of populations (i.e., Tsurugi/Miyazaki) as another fixed factor to avoid complicating the model structure (note that we could not include the study population as a random factor due to the small number of random-effect levels, e.g., Bolker et al. 2009 suggests “>5–6 random-effect levels per random effect” for properly estimating among-group variance). We also conducted analyses of “residual” components of feather coloration controlling for pigment contents, in which, from observed feather coloration, the component explained by the amount of each pigment content was subtracted using the coefficients of the abovementioned models. If the effect of UV exposure on feather coloration reflects on particular pigment content (e.g., protoporphyrin IX), the residual component controlling for the pigment content (but not those for others) would no longer be explained by UV treatment and its interaction with age. Additionally, to further test the effect of pigmentation on feather coloration, we examined difference in the feather coloration between two samples from the same individuals (i.e., one used for control and the other used for UV treatment) in relation to the corresponding difference in pigmentation. In this analysis with a small sample size (*n* = 18), we used a robust linear model (function lmrob in the R package robustbase) to minimize the influence of outliers, as before (Arai et al. 2017). Lastly, we compared the pigment contents of feathers collected from molting adults and those collected from adults and young during the breeding season, to test the effect of time elapsed after feather emergence. All data analyses were performed using the R statistical package (version 4.1.0; R Core Team 2021).

## Results

### Pigmentation per feather

We could quantify PTCA, i.e., an index of eumelanin, TTCA, i.e., an index of pheomelanin, and protoporphyrin IX contents of 38 feather samples. In these samples, the throat feathers of barn swallows contained 13.65 ± 6.96 ng/feather (mean ± SD) PTCA (range: 3.5–26.9 ng/feather), and 13.44 ± 11.22 ng/feather (mean ± SD) TTCA (range: 2.3–56.5 ng/feather). We also detected protoporphyrin IX pigments (mean ± SD = 67.10 ± 97.42 pg/feather, range: 0–403 pg/feather), and its amount was on average 734 times less than TTCA (range: 14–3786 times, excluding two individuals with no detectable protoporphyrin IX content). We could not estimate the amount of protoporphyrin IX pigments in two feather samples from two different individuals, and thus the following analyses of pigmentation consisted of 36 samples from the remaining 18 individuals (note that, as we adopted a “paired” experimental design, each pair of samples should consist of one in control and the other in the UV treatment group). Although we also attempted to analyze coproporphyrin III pigment, substantial amounts of it were not detected in the current samples (i.e., only traces of this pigment were detected in HPLC analysis), preventing further analyses of this pigment. Detection of porphyrin pigments was further reinforced by the fluorescent red coloration of feathers under UV light (Fig. S2).

### UV, coloration, and pigmentation

The interaction term between age and treatment explained the coloration of throat feathers, as measured by hue, saturation, and brightness (Table 1 left column). Feathers exposed to UV light showed significantly higher hue (i.e., less red), saturation, and brightness than control feathers even in adults, as shown by the coefficients of “Treatment” in Table 1 left column, but the effect was particularly strong in young (Fig. 1), as shown by the significant positive interaction terms in Table 1 left column. A similar pattern was found when we analyzed feather color change before and after the experiment (Table 1 right column; Fig. 2).

**Table 1.**
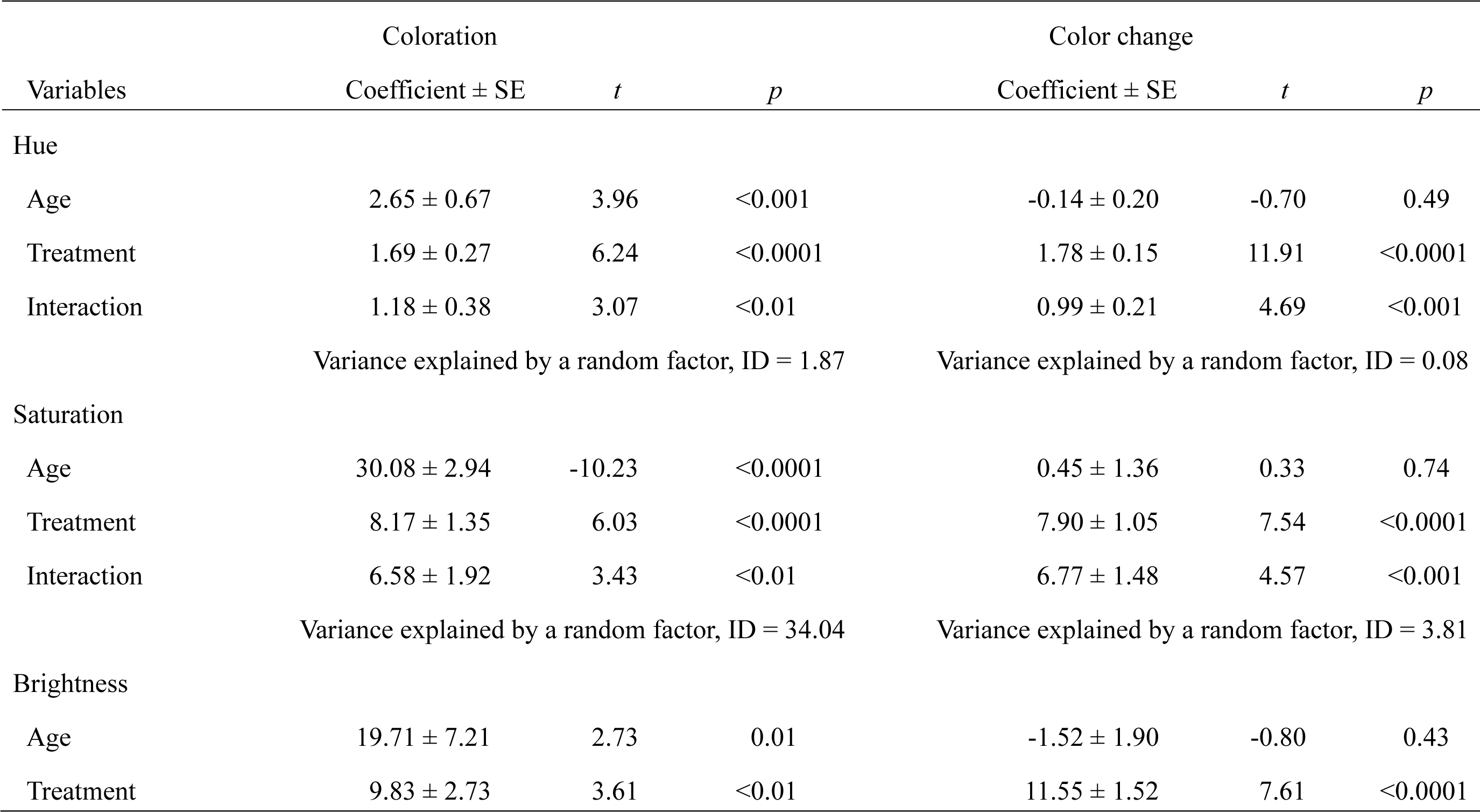
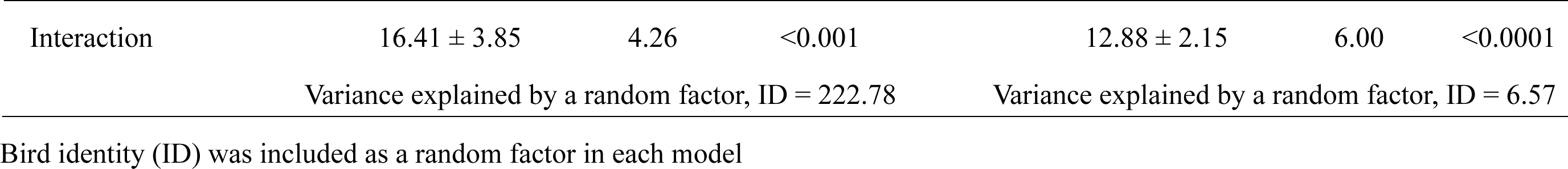
Linear mixed-effects model explaining coloration (left column) and color change (right column) of the throat feathers in relation to age and treatment in the barn swallow (*n_adult,control_* = 10, *n_young,control_* = 10, *n_adult,UV_* = 10, *n_young,UV_* = 10, *n_total_* = 40)

**Figure 1.**
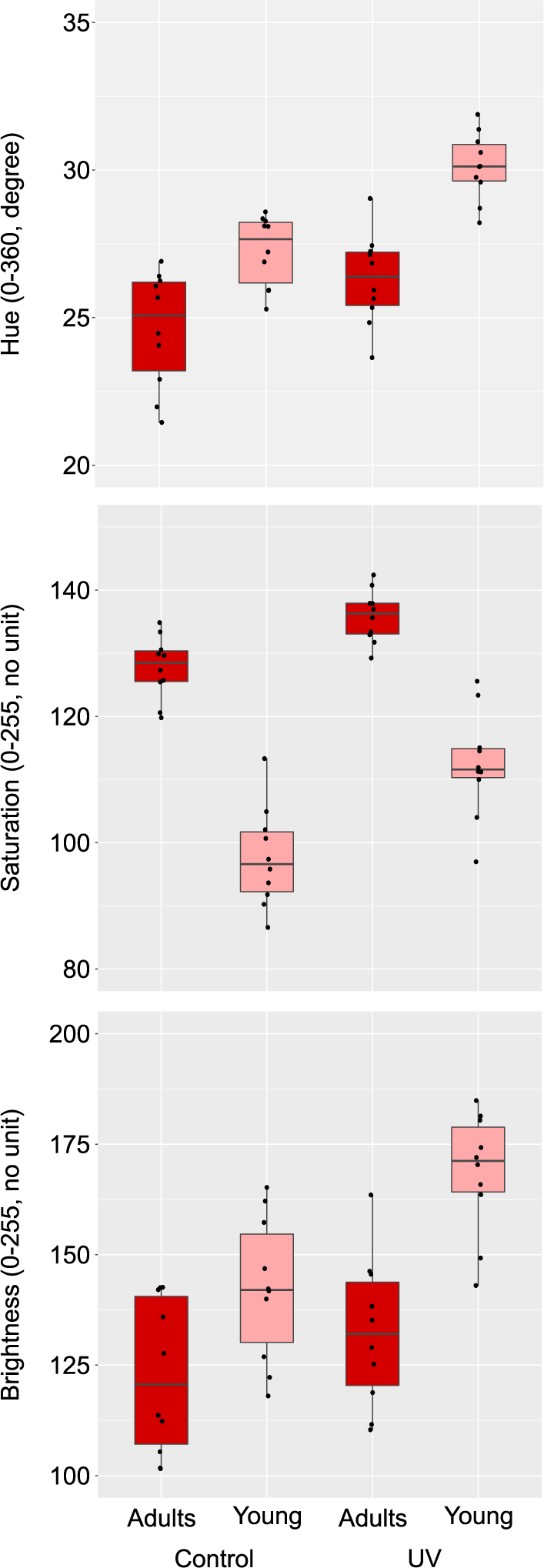
Boxplots of coloration of the throat feathers in relation to age and treatment in the barn swallow: hue (upper panel); saturation (middle panel); brightness (lower panel). The horizontal bar in each boxplot indicates the median, and the box shows the first and third quartiles of data. The whiskers range from the lowest to the highest data points within the 1.5 × interquartile range of the lower and upper quartiles, respectively. Circles indicate each data point. See Table 1 left column for detailed statistics

**Figure 2.**
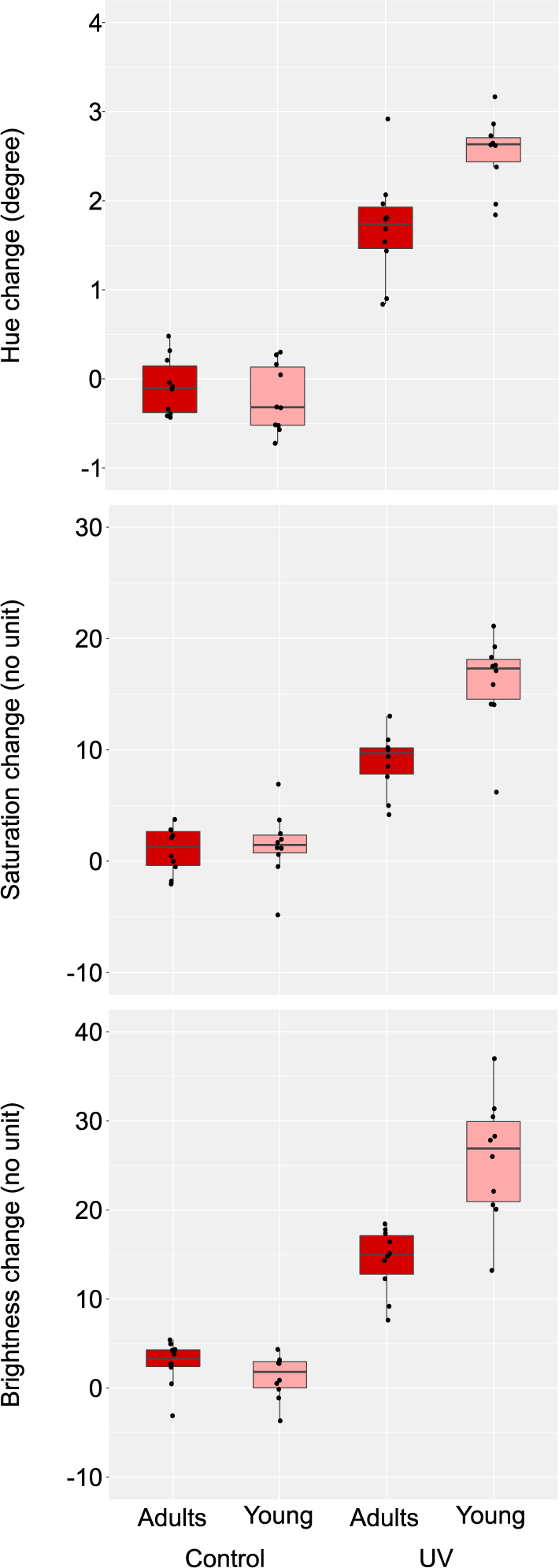
Boxplots of color changes (value after experiment minus value before experiment) of the throat feathers in relation to age and treatment in the barn swallow: hue change (upper panel); saturation change (middle panel); brightness change (lower panel). The horizontal bar in each boxplot indicates the median, and the box shows the first and third quartiles of data. The whiskers range from the lowest to the highest data points within the 1.5 × interquartile range of the lower and upper quartiles, respectively. Circles indicate each data point. See Table 1 right column for detailed statistics

Likewise, the interaction term between age and treatment explained the protoporphyrin IX content of throat feathers (Table 2). Feathers exposed to UV light contained significantly lower amounts of protoporphyrin IX pigments than control feathers, even in adults (ca. 1/3, deduced from Table 2), but again, this effect was particularly strong in young (ca. 1/20, deduced from Table 2; Fig. 3). In contrast, the interaction term between age and treatment was far from significant in PTCA, an index of eumelanin, and TTCA, an index of pheomelanin (Table 2). In these models, treatment remained nonsignificant after the nonsignificant interaction terms were excluded from the models (*p* > 0.43; details not shown).

**Table 2.** Linear mixed-effects model explaining pigmentation of the throat feathers in relation to age and treatment in the barn swallow (*n_adult,control_* = 9, *n_young,control_* = 9, *n_adult,UV_*

| Variables | Coefficient ± SE | *t* | *p* |
| --- | --- | --- | --- |
| Protoporphyrin IX |  |  |  |
| Age | 0.95 ± 0.40 | 2.35 | 0.03 |
| Treatment | -1.22 ± 0.30 | -4.02 | <0.001 |
| Interaction | -1.80 ± 0.43 | -4.20 | <0.001 |
| Variance explained by a random factor, ID = 0.32 |  |  |  |
| PTCA |  |  |  |
| Age | -0.10 ± 0.26 | -0.38 | 0.71 |
| Treatment | -0.01 ± 0.19 | -0.04 | 0.97 |
| Interaction | -0.03 ± 0.27 | -0.11 | 0.91 |
| Variance explained by a random factor, ID = 0.14 |  |  |  |
| TTCA |  |  |  |
| Age | -1.00 ± 0.21 | -4.66 | <0.0001 |
| Treatment | 0.18 ± 0.18 | 0.97 | 0.35 |
| Interaction | -0.15 ± 0.26 | -0.59 | 0.57 |
| Variance explained by a random factor, ID = 0.06 |  |  |  |

Bird identity (ID) was included as a random factor in each model.
Dependent variables were log-transformed before each analysis.
Note that sample sizes were reduced in this analysis because of missing information (see Methods)

**Figure 3.**
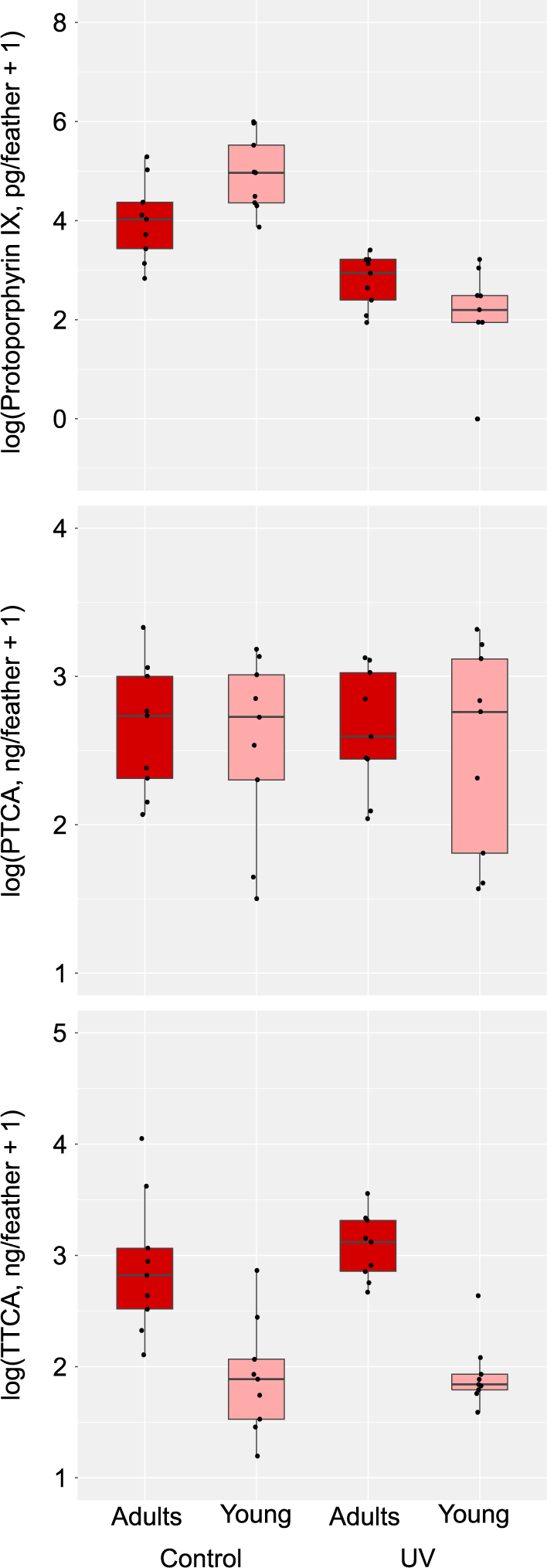
Boxplots of pigmentation of the throat feathers in relation to age and treatment in the barn swallow: protoporphyrin IX (upper panel); PTCA (middle panel); TTCA (lower panel). PTCA and TTCA are indices of eumelanin and pheomelanin, respectively. The horizontal bar in each boxplot indicates the median, and the box shows the first and third quartiles of data. The whiskers range from the lowest to the highest data points within the 1.5 × interquartile range of the lower and upper quartiles, respectively. Circles indicate each data point. See Table 2 for detailed statistics

When we used three pigments as predictor variables in mixed-effects models, protoporphyrin IX explained the variation of the color measurements (Table 3 left column). All three measurements, i.e., hue, saturation, and brightness, decreased with increasing protoporphyrin IX pigments (Fig. 4). Although we also found the significant effect of TTCA, an index of pheomelanin (Table 3 left column), a close inspection of Figure 4 indicates that this was due to between-age differences rather than within-age variation (Fig. 4). In fact, after controlling for the age difference in coloration and pigmentation by subtracting the coefficients of “Age” from measurements of young (see Tables 1 & 2), protoporphyrin IX, but not TTCA, remained significant (protoporphyrin IX: all: *p* < 0.0001; PTCA: all: *p* > 0.36; TTCA: all: *p* > 0.16; details not shown). When we examined feather color change before and after the experiment in relation to the three pigments, protoporphyrin IX but not PTCA or TTCA, explained the color change (Table 3 right column). The more color changed before and after the experiment, the fewer protoporphyrin IX remained in the feathers after the experiment (Fig. 5).

**Table 3.**
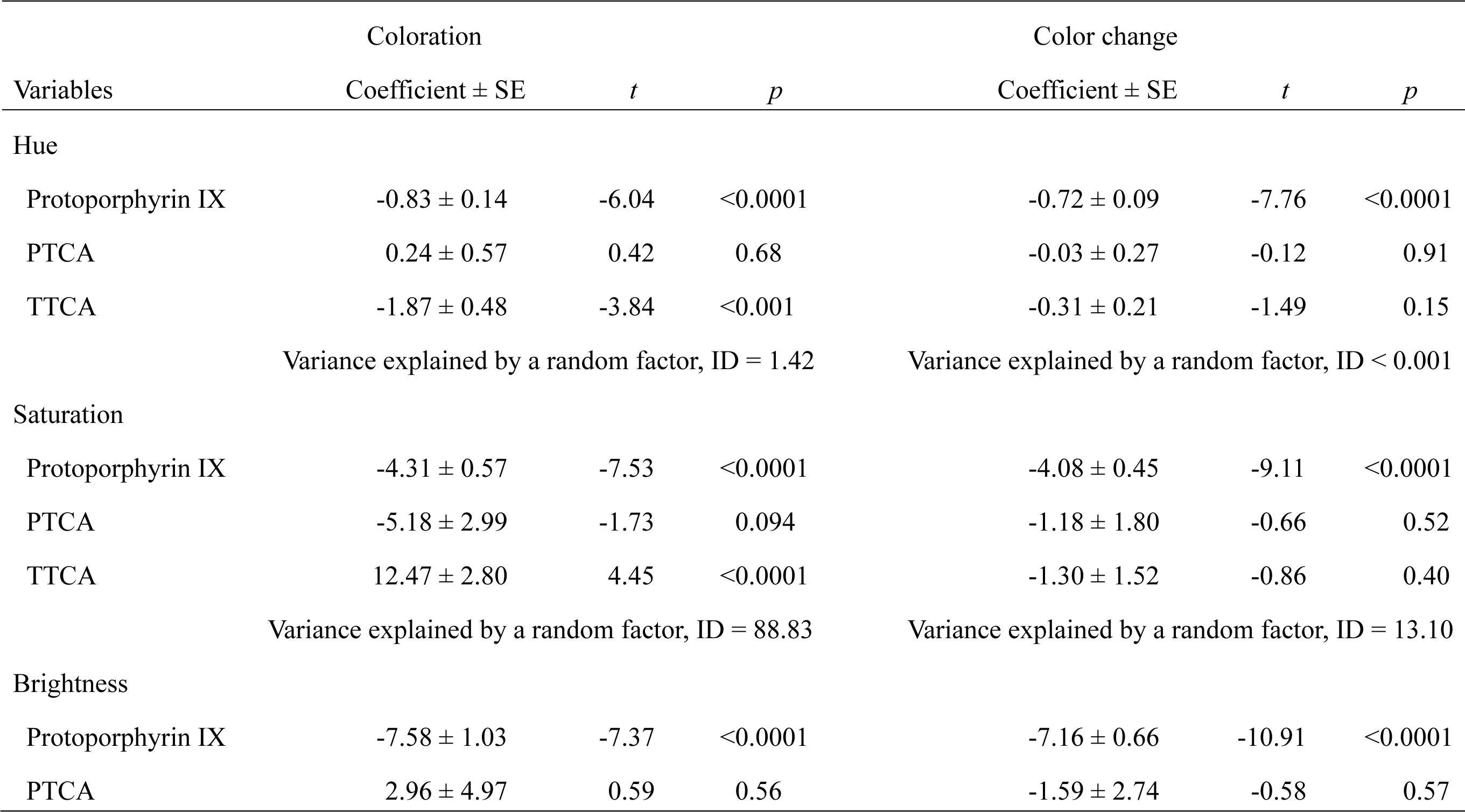
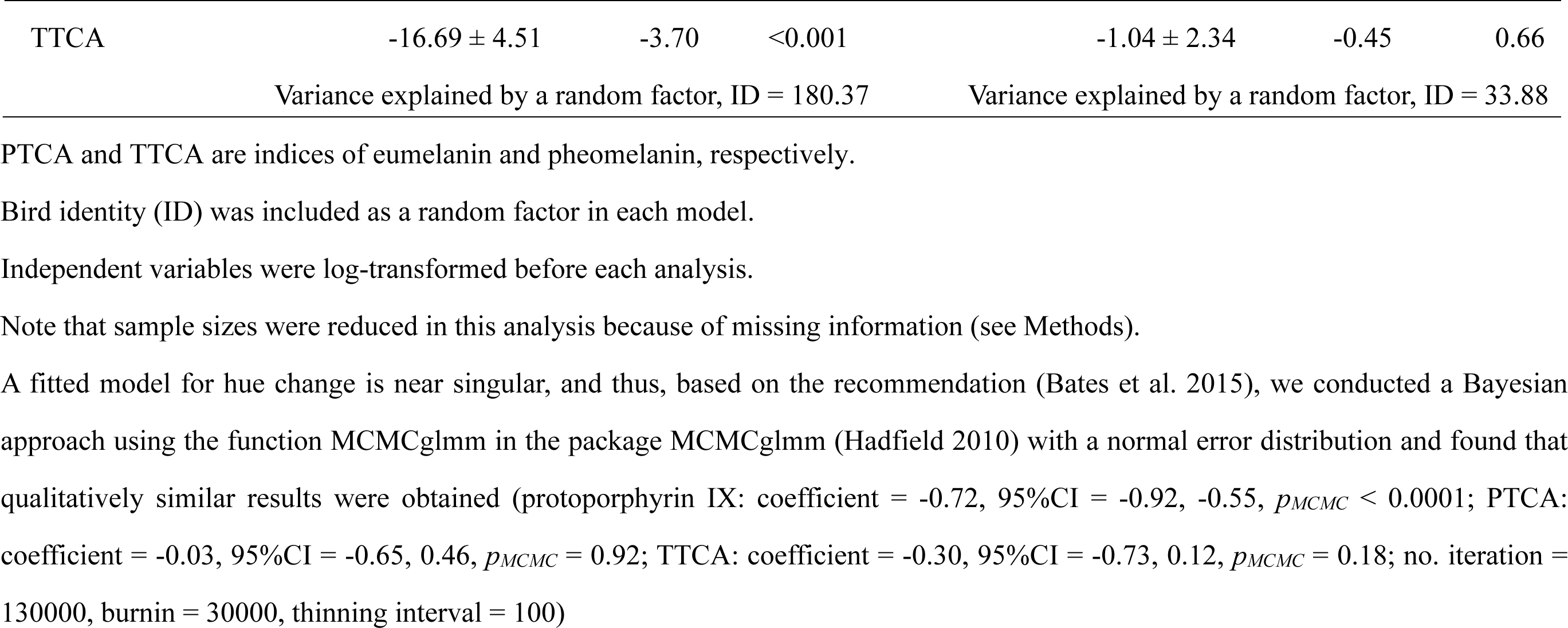
Linear mixed-effects model explaining coloration (left column) and color change (right column) of the throat feathers in relation to pigmentation in the barn swallow (*n_adult,control_* = 9, *n_young,control_* = 9, *n_adult,UV_* = 9, *n_young,UV_* = 9, *n_total_* = 36)

**Figure 4.**
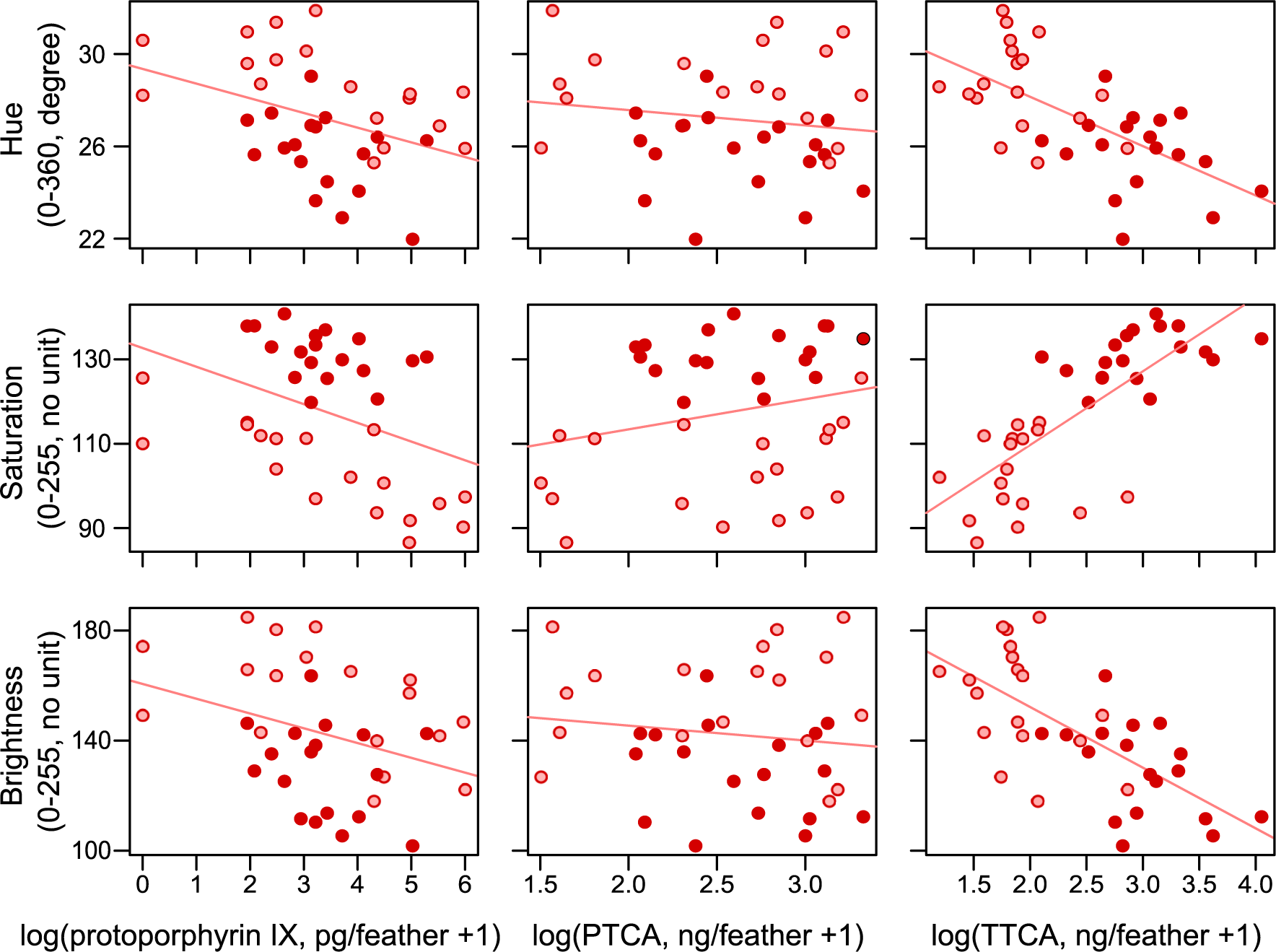
Relationships between color measurements of the throat feathers and pigmentation in the barn swallow: protoporphyrin IX (left column); PTCA (middle column); TTCA (right column). PTCA and TTCA are indices of eumelanin and pheomelanin, respectively. The upper, middle, and bottom rows show hue, saturation, and brightness, respectively. Red and pink circles (dark and light gray, in print) indicate adults and young, respectively. Simple regression lines are depicted for illustrative purpose (See Table 3 left column for detailed statistics)

**Figure 5.**
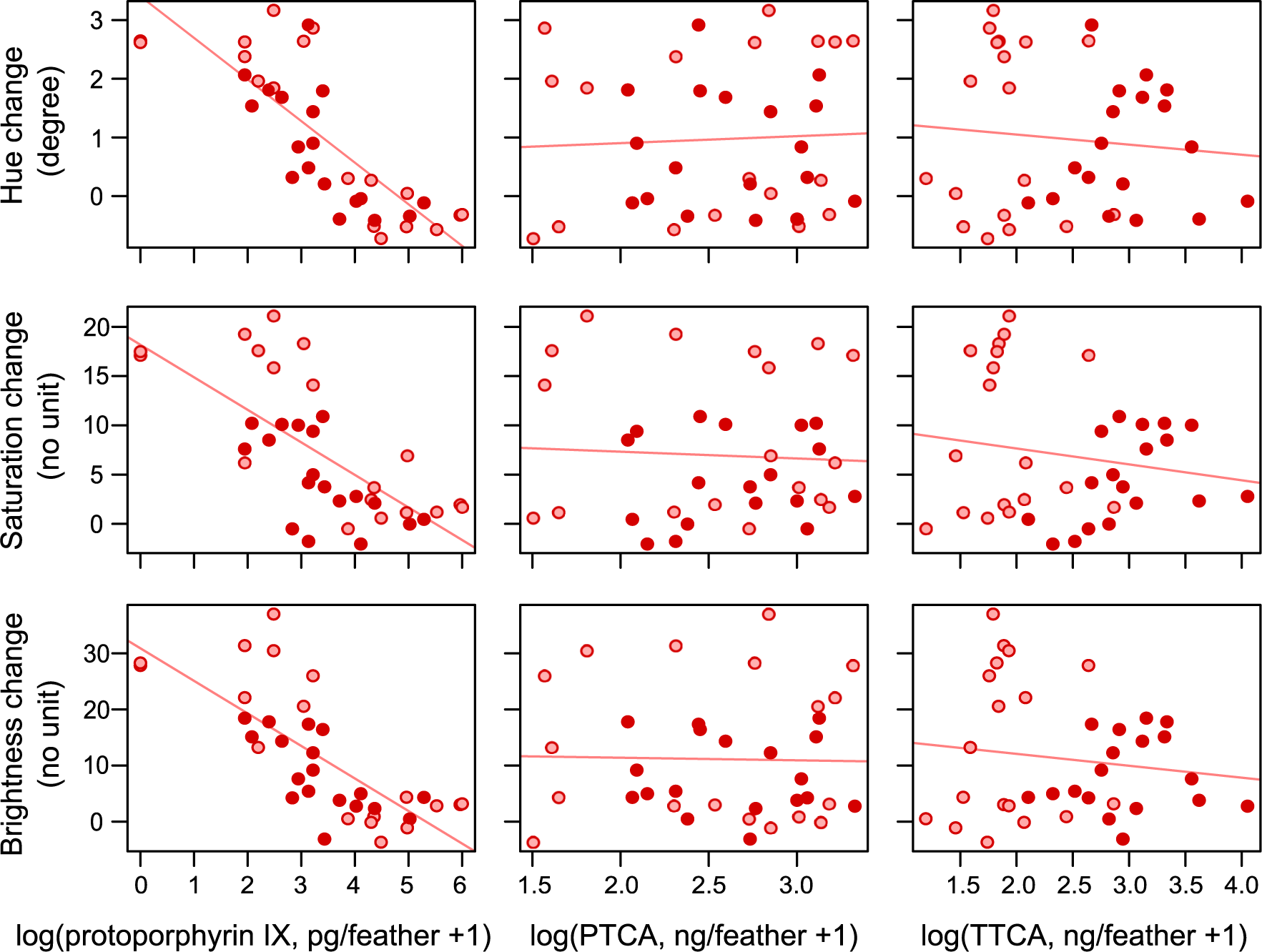
Relationships between color change and pigmentation of the throat feathers in the barn swallow: protoporphyrin IX (left column); PTCA (middle column); TTCA (right column). The upper, middle, and bottom rows show hue, saturation, and brightness, respectively. Red and pink circles (dark and light gray, in print) indicate adults and young, respectively. Simple regression lines are depicted for illustrative purpose (See Table 3 right column for detailed statistics)

After statistically controlling for protoporphyrin IX content, the interaction term between age and treatment on feather coloration and its color change did not remain significant (Tables S3 & S4). This was not the case when we statistically controlled for PTCA or TTCA (i.e., interaction terms remained significant: *p* < 0.05; details not shown). An alternative approach to study the linkage between coloration and pigmentation is to examine the difference in pigmentation between the two treatments collected from the same individuals (i.e., one in the UV treatment group and the other in the Control; see Methods) in relation to the corresponding difference in coloration between them, assuming that the effect of UV treatment on coloration reflects that on its pigment content. As predicted, we found that the difference in protoporphyrin IX consistently explained the difference in each of the three coloration measurements, although the analysis of hue difference was marginal (i.e., *p* = 0.05; Table S2). The difference in the color measurements between the two treatments increased with the increasing reduction of protoporphyrin IX pigments in the UV treatment group compared with controls (Fig. S3).

### Newly molted feathers

When comparing protoporphyrin IX pigments of feathers collected from molting adults in winter (Fig. S1) with those collected from adults and young during the breeding season, we found that the amount of protoporphyrin IX content of throat feathers collected from molting adults was comparable to that of throat feathers collected from young during the breeding season but was significantly higher than that of throat feathers collected from adults during the breeding season (ca. five times, deduced from Table 4; Fig. 6). There were no detectable differences in the amount of eumelanin and pheomelanin content of throat feathers between molting adults and the other two groups, although the difference in the amount of pheomelanin content between throat feathers collected from young and molting adults was marginal (Table 4; Fig. 6).

**Table 4.** Linear mixed-effects model explaining pigmentation of the throat feathers in the barn swallow including newly molted adults (*n_post-molting adult_* = 3, *n_breeding adult_* = 10, *n_young_* = 9, *n_total_* = 22)

| Variables | Coefficient ± SE | *t* | *p* |
| --- | --- | --- | --- |
| Protoporphyrin IX |  |  |  |
| Difference to breeding adults | $-1.54 \pm 0.49$ | -3.16 | <0.01 |
| Difference to young | $-0.57 \pm 0.49$ | -1.16 | 0.26 |
| PTCA |  |  |  |
| Difference to breeding adults | $-0.08 \pm 0.39$ | -0.20 | 0.84 |
| Difference to young | $-0.08 \pm 0.40$ | -0.20 | 0.85 |
| TTCA |  |  |  |
| Difference to breeding adults | $0.24 \pm 0.38$ | 0.62 | 0.54 |
| Difference to young | $-0.69 \pm 0.39$ | -1.79 | 0.090 |

Dependent variables were log-transformed before each analysis

**Figure 6.**
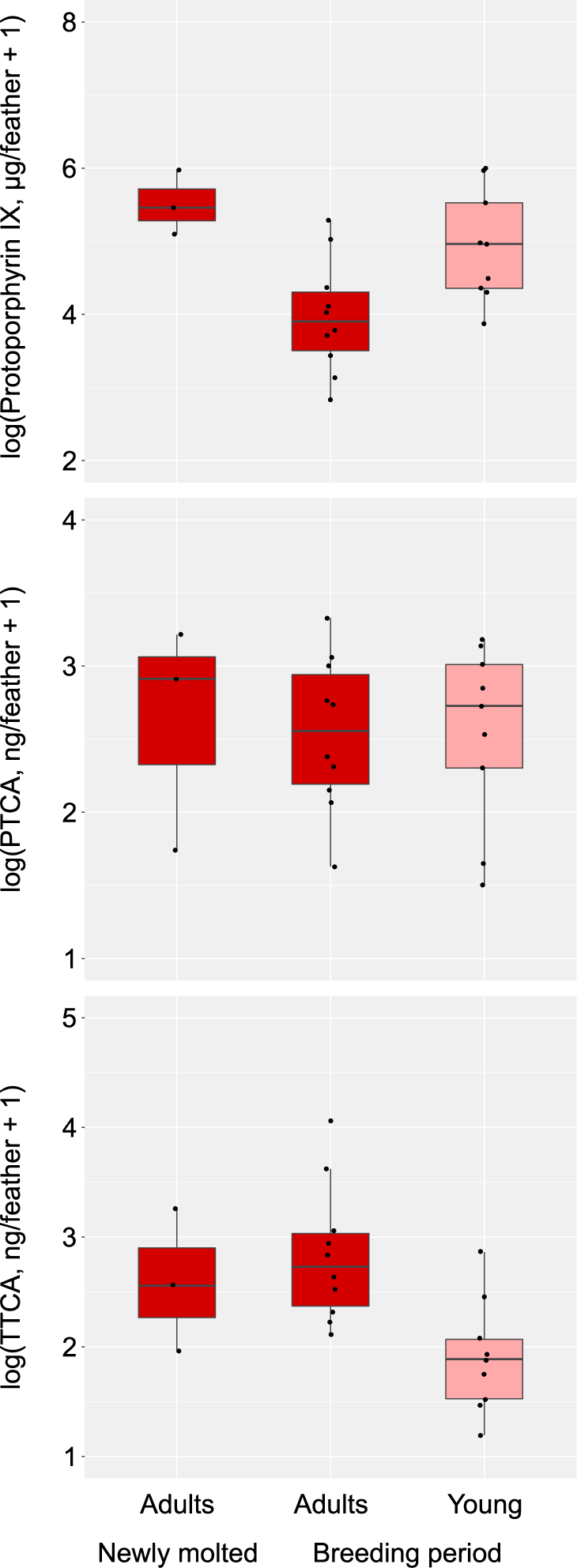
Boxplots of pigmentation of the throat feathers in relation to molting adults and adults and young captured during the breeding season in the barn swallow: protoporphyrin IX (upper column); PTCA (middle column); TTCA (lower column). PTCA and TTCA are indices of eumelanin and pheomelanin, respectively. The horizontal bar in each boxplot indicates the median, and the box shows the first and third quartiles of data. The whiskers range from the lowest to the highest data points within the 1.5 × interquartile range of the lower and upper quartiles, respectively. See Table 4 for detailed statistics

## Discussion

The main finding of the current study is that protoporphyrin IX pigments, seemingly an “uncommon pigment” (McGraw 2006c), explained the pheomelanin-based feather coloration of barn swallows and its age-dependent color change under the exposure of UV light. This finding is understandable due to the wide range of protoporphyrin IX content of the feathers but is surprising because the protoporphyrin IX content of throat feathers was, on average, >500 times less than eumelanin and pheomelanin contents in the barn swallow. The observed porphyrin content is low even when compared with other species, such as the barn owl *Tyto alba*, in which protoporphyrin IX content was one tenth of pheomelanin pigments (Roulin et al. 2008). Previous studies often focused on the predominant pigments such as carotenoids and melanins, assuming that the evolutionary and ecological features of integument coloration can be attributed to the chemical properties of these predominant pigments (e.g., see McGraw 2006b, Ducrest et al. 2008, Roulin 2016 for reviews on melanin-based feather coloration). However, as was shown here, feather color change, an ecological feature of feather coloration, depends on a minority pigment, and thus the assumption that predominant pigments determine the evolutionary and ecological features of animal coloration is no longer valid. Because feather coloration changed even in adults (Table 1), porphyrin would affect feather coloration of adults as well, although the feather color change of adults was smaller than that of young (Table 1).

A fundamental question about porphyrin is why birds (and some other animals) use porphyrins as colorants in addition to other pigments (e.g., pheomelanin, here). A possible explanation is that, when the use of coloration is confined to a short period after feather emergence, birds use porphyrin to temporally “boost” feather coloration (Galván et al. 2016, 2018). In the barn swallow, nestling feather coloration affects parental food allocation (Romano et al. 2016; Arai et al. unpublished data), which lasts for only a few weeks before the independence of offspring (Turner 2006). Then, for young birds, pigments that can easily be photodegraded (i.e., porphyrin) are preferable to relatively stable pigments (i.e., pheomelanin, here), because retaining conspicuous coloration after independence would incur viability costs (e.g., via increased predatory attack).

A relatively large amount of protoporphyrin IX pigments found in young, in combination with a relatively low amount of pheomelanin pigments (Table 2), at first seem consistent with this explanation. Young swallows might use many porphyrin pigments to rapidly fade their coloration in their open (and thus UV abundant) habitat after its use. Susceptibility to UV light, however, provides an alternative and simpler explanation for this pattern. In contrast with adults, which molt after the breeding season (see Methods), nestlings grow their feathers in the breeding nest just before fledging; therefore, much more protoporphyrin pigments would remain in the feathers at sampling. In fact, even in adults, we found a relatively large amount of protoporphyrin IX in their newly molted feathers (Table 4; Fig. 6), supporting this perspective. Thus, the deposition of a large amount of protoporphyrin in feathers would not be unique to nestlings but be shared with adults. For this reason, we cannot exclude the possibility that the supposedly “adaptive” function of porphyrin in young (see above) might be an exaptation (i.e., secondary use of already evolved traits for other functions: Bergstrom and Dugatkin 2016). Rather, as we found a substantial amount of protoporphyrin IX in breeding adults, adults (rather than young) might evolve unique mechanisms to slow down the photodegradation of porphyrin, which might be beneficial for them to retain their coloration until the end of the breeding period. Although age-dependent depigmentation of protoporphyrin IX (Table 2) is consistent with this perspective, we have no data to test this possibility in the current data set. The ecological function of porphyrin and its evolutionary importance remains to be clarified in future studies, by focusing not only on pigment production but also on its maintenance.

A caveat of the current study is that we focused only on feather color change under ultraviolet light, particularly, what is called, UV-A light (i.e., wavelength: 320–400 nm). We chose UV-A, simply because the stratospheric ozone absorbs most UV-B radiation and thus UV-A light is much more abundant in the environment (e.g., see Japan Meteorological Agency 2010), and because UV-A light penetrates deeper integument layers than UV-B light (and thus we speculated that this could explain the uniform depigmentation of pre-molt feathers). However, feather color change can also occur for other reasons, such as photodegradation via other wavelengths of sunlight including UV-B, feather abrasion, and accumulation of stains (reviewed in Montgomerie 2006). In fact, using the same study species, Arai et al. (2015) showed that, in the wild, the amount of pheomelanin content in throat feathers decreased at a rate of 15–25% per month, and thus these components should also be involved in the wild, together with the photodegradation of porphyrin pigments we demonstrated here (e.g., the amount of porphyrin content decreased at a rate of 30–90% per month, assuming that UV exposure is confined to daytime; Table 2). In addition, because the amount of protoporphyrin IX could not explain all the feather color changes (Table S4), even under the current experimental condition, not only pigmentation but also other factors, such as preen oil, feather microstructures, and their interactions with pigmentation, might be involved in feather color change, which should be studied in the future to understand the whole picture of feather color change.

In summary, we demonstrated that an uncommon pigment, protoporphyrin IX, affected pheomelanin-based coloration and its dynamic color change under UV light. Feather color change, however, would not be the sole ecological feature of feather porphyrin. For example, protoporphyrin IX is a well-known colorant of eggs (reviewed in McGraw 2006c), and its expression is known to correlate with hatching success and female feather coloration, at least in the barn swallow (Corti et al. 2018). Then, it is likely that the porphyrin content of feathers can partially explain the linkage, serving feather coloration as an honest signal of the reproductive value of their clutches. It remains to be clarified how these minority pigments affect the overall evolution and ecology of animal coloration in future studies. In particular, the co-occurrence of porphyrin and pheomelanin, i.e., two similar-colored pigments with different chemical properties (McGraw 2006c), and their inter- and intraspecific variation, would provide a unique study system to understand how particular pigments and their combinations (or possibly displacement) evolved to produce the same kind of coloration.

## Declarations

### Author contribution

MH conceived the ideas; MH and EA designed methodology; MH conducted field survey, collected the data of color measurements, and analyzed the data, and led the writing of the manuscript; EA prepared samples for HPLC analyses; SI and KW conducted HPLC analyses and collected the data of pigmentation. All authors contributed to final draft and gave final approval for publication.

### Data archiving

The datasets supporting this article have been uploaded as part of electronic supplementary material, table S1, which will be uploaded to osf.io after acceptance.

### Ethical approval

The permits for the current study including capturing were provided by Ishikawa Prefecture (#20249) and Miyazaki Prefecture (#100) and Ishikawa Prefectural University (#R4-14-2, #R4-14-23), following the Wildlife Protection and Hunting Management Law.

## Acknowledgments

We are grateful to the residents of Tsurugi Town and Miyazaki City for their kind support and assistance. We also thank Dr. Ichiro Tayasu, and Dr. Shumpei Kitamura and his laboratory members in Ishikawa Prefectural University.

---

**Table S1.**
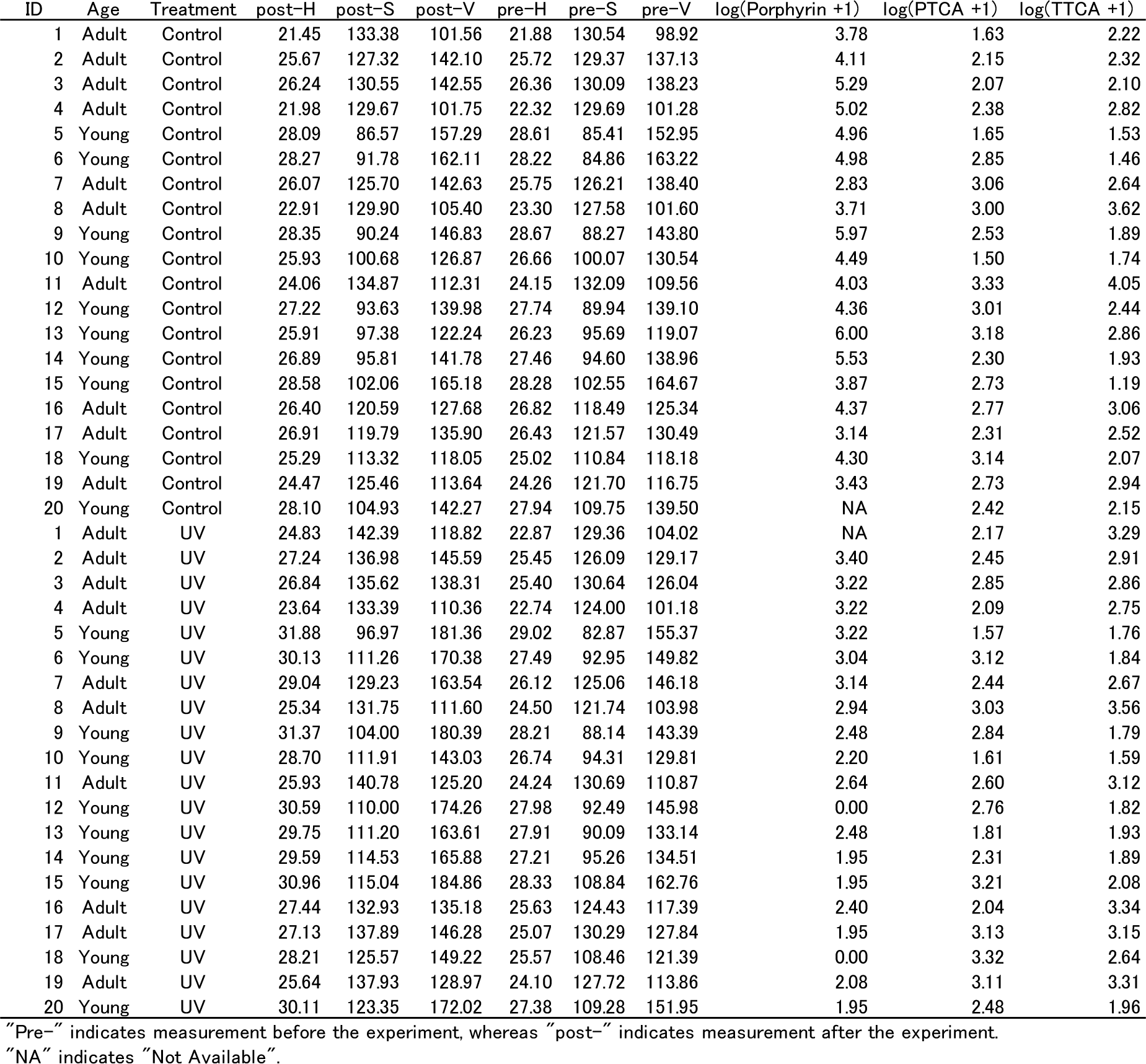
Dataset of the current study

| ID | Age | Treatment | post-H | post-S | post-V | pre-H | pre-S | pre-V | log(Porphyrin +1) | log(PTCA +1) | log(TTCA +1) |
| --- | --- | --- | --- | --- | --- | --- | --- | --- | --- | --- | --- |
| 1 | Adult | Control | 21.45 | 133.38 | 101.56 | 21.88 | 130.54 | 98.92 | 3.78 | 1.63 | 2.22 |
| 2 | Adult | Control | 25.67 | 127.32 | 142.10 | 25.72 | 129.37 | 137.13 | 4.11 | 2.15 | 2.32 |
| 3 | Adult | Control | 26.24 | 130.55 | 142.55 | 26.36 | 130.09 | 138.23 | 5.29 | 2.07 | 2.10 |
| 4 | Adult | Control | 21.98 | 129.67 | 101.75 | 22.32 | 129.69 | 101.28 | 5.02 | 2.38 | 2.82 |
| 5 | Young | Control | 28.09 | 86.57 | 157.29 | 28.61 | 85.41 | 152.95 | 4.96 | 1.65 | 1.53 |
| 6 | Young | Control | 28.27 | 91.78 | 162.11 | 28.22 | 84.86 | 163.22 | 4.98 | 2.85 | 1.46 |
| 7 | Adult | Control | 26.07 | 125.70 | 142.63 | 25.75 | 126.21 | 138.40 | 2.83 | 3.06 | 2.64 |
| 8 | Adult | Control | 22.91 | 129.90 | 105.40 | 23.30 | 127.58 | 101.60 | 3.71 | 3.00 | 3.62 |
| 9 | Young | Control | 28.35 | 90.24 | 146.83 | 28.67 | 88.27 | 143.80 | 5.97 | 2.53 | 1.89 |
| 10 | Young | Control | 25.93 | 100.68 | 126.87 | 26.66 | 100.07 | 130.54 | 4.49 | 1.50 | 1.74 |
| 11 | Adult | Control | 24.06 | 134.87 | 112.31 | 24.15 | 132.09 | 109.56 | 4.03 | 3.33 | 4.05 |
| 12 | Young | Control | 27.22 | 93.63 | 139.98 | 27.74 | 89.94 | 139.10 | 4.36 | 3.01 | 2.44 |
| 13 | Young | Control | 25.91 | 97.38 | 122.24 | 26.23 | 95.69 | 119.07 | 6.00 | 3.18 | 2.86 |
| 14 | Young | Control | 26.89 | 95.81 | 141.78 | 27.46 | 94.60 | 138.96 | 5.53 | 2.30 | 1.93 |
| 15 | Young | Control | 28.58 | 102.06 | 165.18 | 28.28 | 102.55 | 164.67 | 3.87 | 2.73 | 1.19 |
| 16 | Adult | Control | 26.40 | 120.59 | 127.68 | 26.82 | 118.49 | 125.34 | 4.37 | 2.77 | 3.06 |
| 17 | Adult | Control | 26.91 | 119.79 | 135.90 | 26.43 | 121.57 | 130.49 | 3.14 | 2.31 | 2.52 |
| 18 | Young | Control | 25.29 | 113.32 | 118.05 | 25.02 | 110.84 | 118.18 | 4.30 | 3.14 | 2.07 |
| 19 | Adult | Control | 24.47 | 125.46 | 113.64 | 24.26 | 121.70 | 116.75 | 3.43 | 2.73 | 2.94 |
| 20 | Young | Control | 28.10 | 104.93 | 142.27 | 27.94 | 109.75 | 139.50 | NA | 2.42 | 2.15 |
| 1 | Adult | UV | 24.83 | 142.39 | 118.82 | 22.87 | 129.36 | 104.02 | NA | 2.17 | 3.29 |
| 2 | Adult | UV | 27.24 | 136.98 | 145.59 | 25.45 | 126.09 | 129.17 | 3.40 | 2.45 | 2.91 |
| 3 | Adult | UV | 26.84 | 135.62 | 138.31 | 25.40 | 130.64 | 126.04 | 3.22 | 2.85 | 2.86 |
| 4 | Adult | UV | 23.64 | 133.39 | 110.36 | 22.74 | 124.00 | 101.18 | 3.22 | 2.09 | 2.75 |
| 5 | Young | UV | 31.88 | 96.97 | 181.36 | 29.02 | 82.87 | 155.37 | 3.22 | 1.57 | 1.76 |
| 6 | Young | UV | 30.13 | 111.26 | 170.38 | 27.49 | 92.95 | 149.82 | 3.04 | 3.12 | 1.84 |
| 7 | Adult | UV | 29.04 | 129.23 | 163.54 | 26.12 | 125.06 | 146.18 | 3.14 | 2.44 | 2.67 |
| 8 | Adult | UV | 25.34 | 131.75 | 111.60 | 24.50 | 121.74 | 103.98 | 2.94 | 3.03 | 3.56 |
| 9 | Young | UV | 31.37 | 104.00 | 180.39 | 28.21 | 88.14 | 143.39 | 2.48 | 2.84 | 1.79 |
| 10 | Young | UV | 28.70 | 111.91 | 143.03 | 26.74 | 94.31 | 129.81 | 2.20 | 1.61 | 1.59 |
| 11 | Adult | UV | 25.93 | 140.78 | 125.20 | 24.24 | 130.69 | 110.87 | 2.64 | 2.60 | 3.12 |
| 12 | Young | UV | 30.59 | 110.00 | 174.26 | 27.98 | 92.49 | 145.98 | 0.00 | 2.76 | 1.82 |
| 13 | Young | UV | 29.75 | 111.20 | 163.61 | 27.91 | 90.09 | 133.14 | 2.48 | 1.81 | 1.93 |
| 14 | Young | UV | 29.59 | 114.53 | 165.88 | 27.21 | 95.26 | 134.51 | 1.95 | 2.31 | 1.89 |
| 15 | Young | UV | 30.96 | 115.04 | 184.86 | 28.33 | 108.84 | 162.76 | 1.95 | 3.21 | 2.08 |
| 16 | Adult | UV | 27.44 | 132.93 | 135.18 | 25.63 | 124.43 | 117.39 | 2.40 | 2.04 | 3.34 |
| 17 | Adult | UV | 27.13 | 137.89 | 146.28 | 25.07 | 130.29 | 127.84 | 1.95 | 3.13 | 3.15 |
| 18 | Young | UV | 28.21 | 125.57 | 149.22 | 25.57 | 108.46 | 121.39 | 0.00 | 3.32 | 2.64 |
| 19 | Adult | UV | 25.64 | 137.93 | 128.97 | 24.10 | 127.72 | 113.86 | 2.08 | 3.11 | 3.31 |
| 20 | Young | UV | 30.11 | 123.35 | 172.02 | 27.38 | 109.28 | 151.95 | 1.95 | 2.48 | 1.96 |

"Pre-" indicates measurement before the experiment, whereas "post-" indicates measurement after the experiment.
"NA" indicates "Not Available".

**Table S2.** Linear mixed-effects model explaining the color difference of the throat feathers between treatments in relation to pigmentation difference between treatments in the barn swallow (*n_adult_* = 9, *n_young_* = 9, *n_total_* = 18)

| Variables | Coefficient ± SE | *t* | *p* |
| --- | --- | --- | --- |
| Hue difference |  |  |  |
| Protoporphyrin IX difference | $-0.32 \pm 0.15$ | -2.10 | 0.0545 |
| PTCA difference | $-0.69 \pm 0.92$ | -0.75 | 0.47 |
| TTCA difference | $-0.24 \pm 0.92$ | -0.26 | 0.80 |
| Saturation |  |  |  |
| Protoporphyrin IX difference | $-2.56 \pm 0.93$ | -2.74 | 0.016 |
| PTCA difference | $0.96 \pm 2.88$ | 0.33 | 0.74 |
| TTCA difference | $1.72 \pm 2.56$ | 0.67 | 0.52 |
| Brightness |  |  |  |
| Protoporphyrin IX difference | $-6.27 \pm 2.01$ | -3.12 | <0.01 |
| PTCA difference | $-5.92 \pm 12.29$ | -0.48 | 0.64 |
| TTCA difference | $-1.33 \pm 9.09$ | -0.15 | 0.89 |

PTCA and TTCA are indices of eumelanin and pheomelanin, respectively.
Note that, due to the small sample sizes in these analyses, in which small deviation from normal distribution affects results, we here used robust linear models (function `lmrob` in the R package `robustbase`) as before (Arai et al. 2017).
Independent variables were log-transformed before each analysis

**Table S3.**
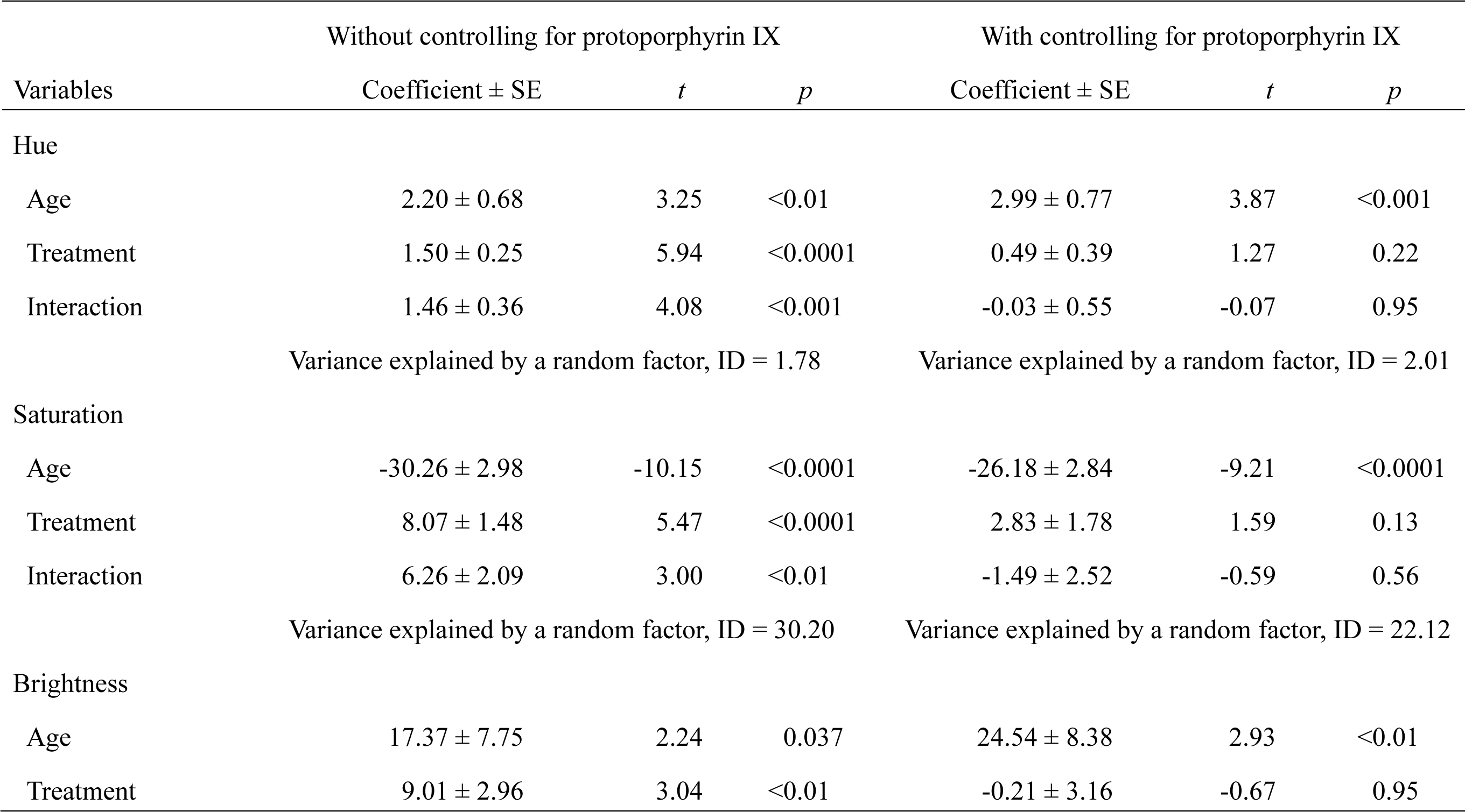
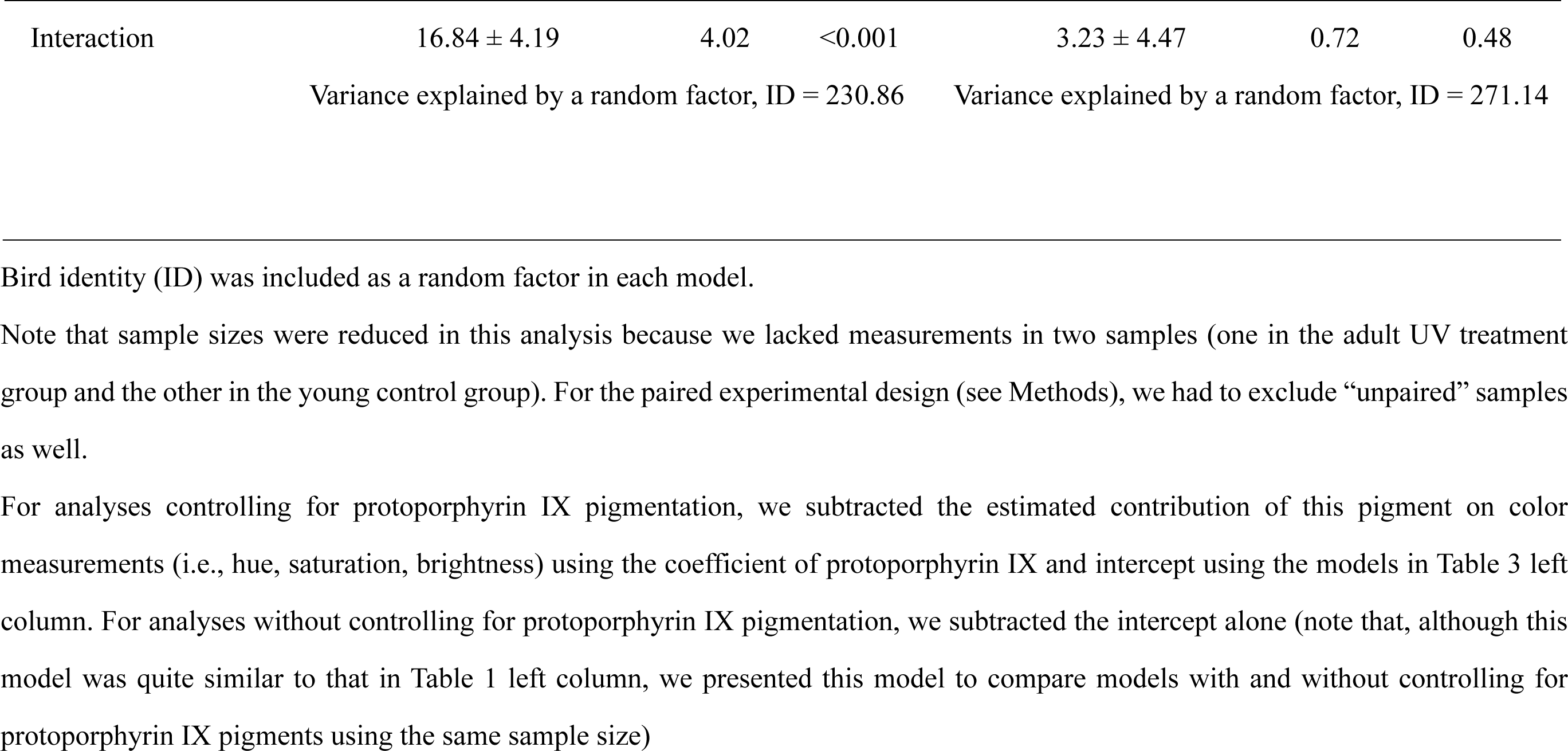
Linear mixed-effects model explaining coloration of the throat feathers in relation to age and treatment with and without controlling for protoporphyrin IX in the barn swallow (*n_adult,control_* = 9, *n_young,control_* = 9, *n_adult,UV_* = 9, *n_young,UV_* = 9, *n_total_* = 36)

|  | Without controlling for protoporphyrin IX |  |  | With controlling for protoporphyrin IX |  |  |
| --- | --- | --- | --- | --- | --- | --- |
| Variables | Coefficient ± SE | *t* | *p* | Coefficient ± SE | *t* | *p* |
| Hue |  |  |  |  |  |  |
| Age | 2.20 ± 0.68 | 3.25 | <0.01 | 2.99 ± 0.77 | 3.87 | <0.001 |
| Treatment | 1.50 ± 0.25 | 5.94 | <0.0001 | 0.49 ± 0.39 | 1.27 | 0.22 |
| Interaction | 1.46 ± 0.36 | 4.08 | <0.001 | -0.03 ± 0.55 | -0.07 | 0.95 |
|  | Variance explained by a random factor, ID = 1.78 |  |  | Variance explained by a random factor, ID = 2.01 |  |  |
| Saturation |  |  |  |  |  |  |
| Age | -30.26 ± 2.98 | -10.15 | <0.0001 | -26.18 ± 2.84 | -9.21 | <0.0001 |
| Treatment | 8.07 ± 1.48 | 5.47 | <0.0001 | 2.83 ± 1.78 | 1.59 | 0.13 |
| Interaction | 6.26 ± 2.09 | 3.00 | <0.01 | -1.49 ± 2.52 | -0.59 | 0.56 |
|  | Variance explained by a random factor, ID = 30.20 |  |  | Variance explained by a random factor, ID = 22.12 |  |  |
| Brightness |  |  |  |  |  |  |
| Age | 17.37 ± 7.75 | 2.24 | 0.037 | 24.54 ± 8.38 | 2.93 | <0.01 |
| Treatment | 9.01 ± 2.96 | 3.04 | <0.01 | -0.21 ± 3.16 | -0.67 | 0.95 |
| Interaction | 16.84 ± 4.19 | 4.02 | <0.001 | 3.23 ± 4.47 | 0.72 | 0.48 |
|  | Variance explained by a random factor, ID = 230.86 |  |  | Variance explained by a random factor, ID = 271.14 |  |  |

Bird identity (ID) was included as a random factor in each model.
Note that sample sizes were reduced in this analysis because we lacked measurements in two samples (one in the adult UV treatment group and the other in the young control group). For the paired experimental design (see Methods), we had to exclude “unpaired” samples as well.

**Table S4.**
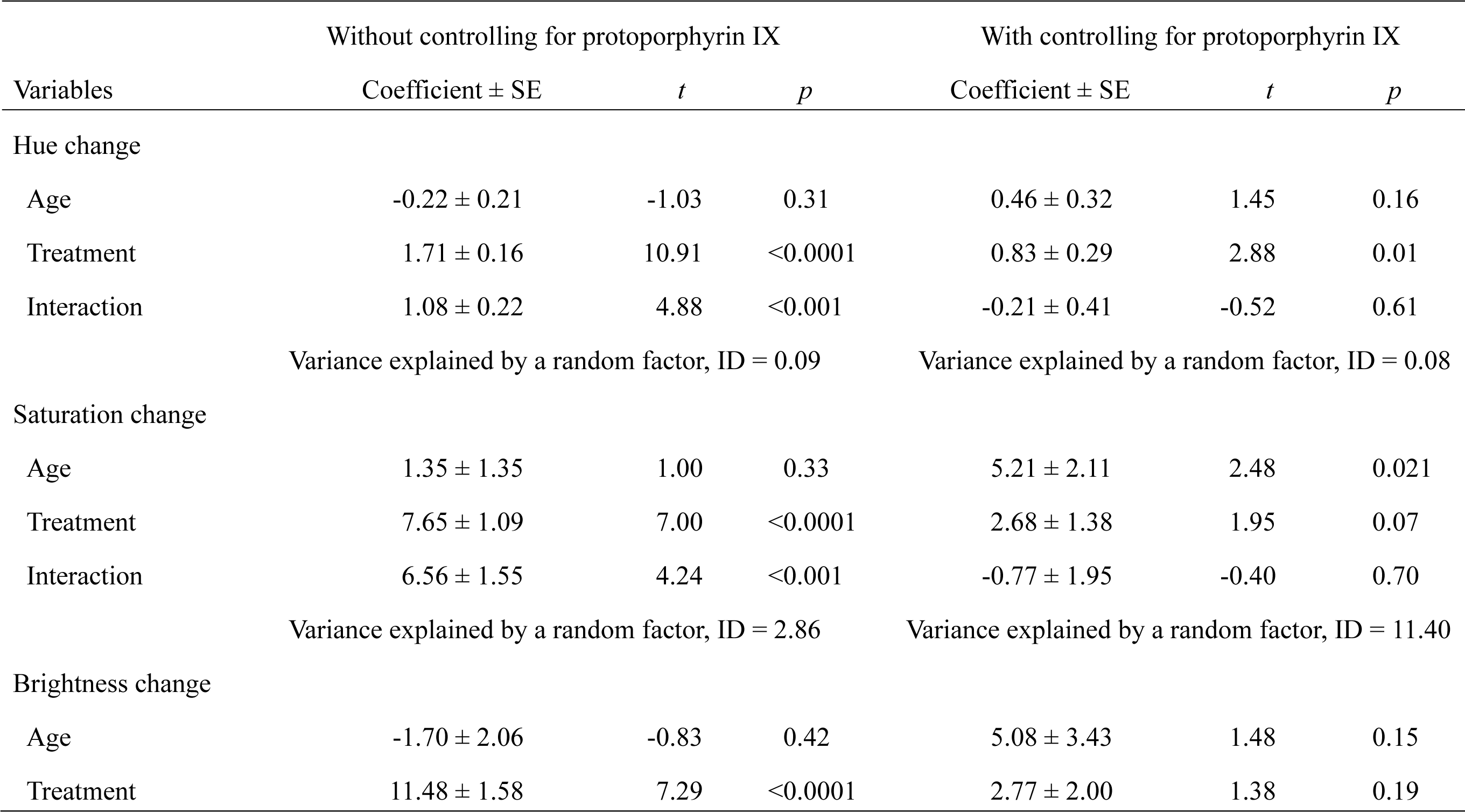
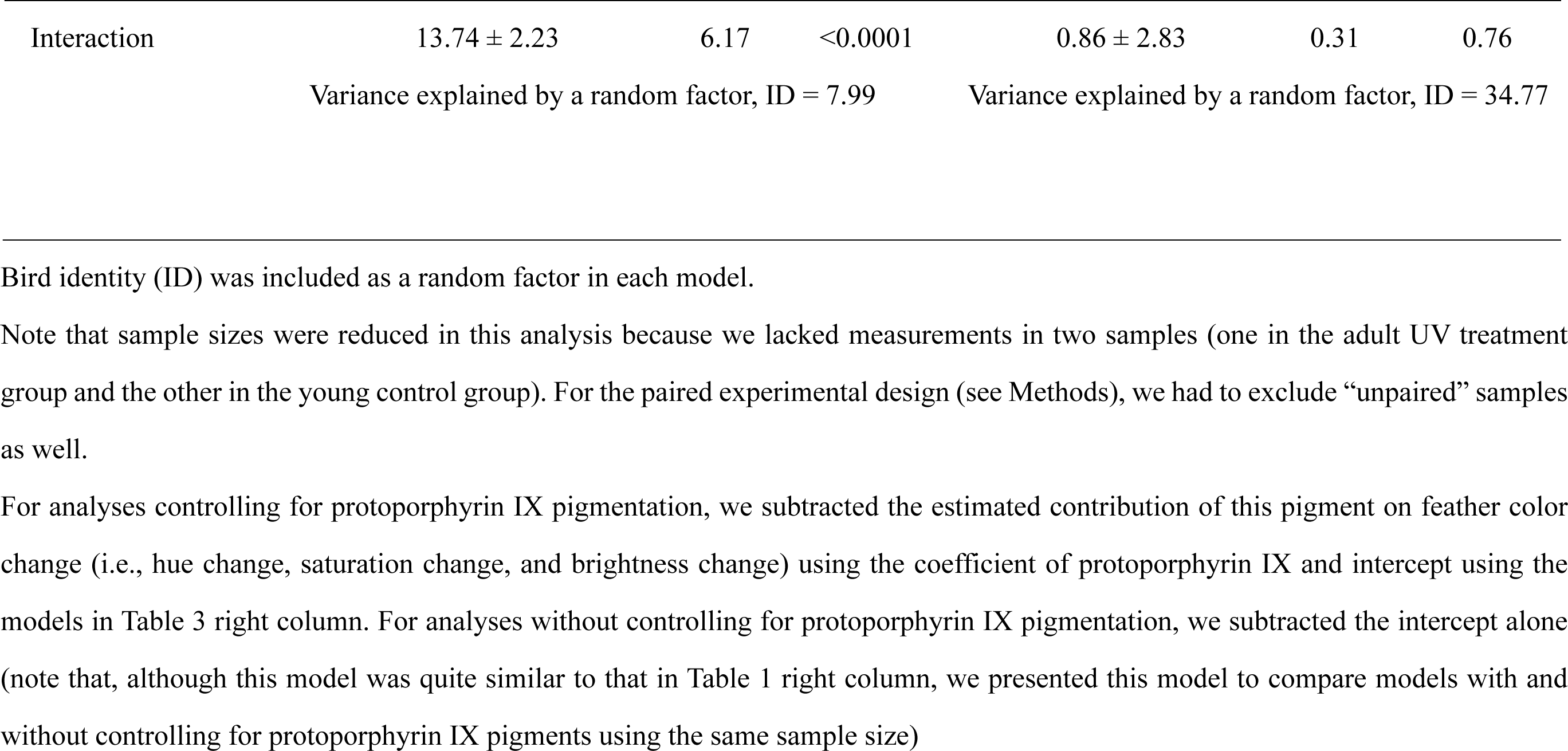
Linear mixed-effects model explaining plumage color change of the throat feathers in relation to age and treatment in the barn swallow with and without controlling for protoporphyrin IX (*n_adult,control_* = 9, *n_young,control_* = 9, *n_adult,UV_* = 9, *n_young,UV_* = 9, *n_total_* = 36)

|  | Without controlling for protoporphyrin IX |  |  | With controlling for protoporphyrin IX |  |  |
| --- | --- | --- | --- | --- | --- | --- |
| Variables | Coefficient ± SE | *t* | *p* | Coefficient ± SE | *t* | *p* |
| Hue change |  |  |  |  |  |  |
| Age | -0.22 ± 0.21 | -1.03 | 0.31 | 0.46 ± 0.32 | 1.45 | 0.16 |
| Treatment | 1.71 ± 0.16 | 10.91 | <0.0001 | 0.83 ± 0.29 | 2.88 | 0.01 |
| Interaction | 1.08 ± 0.22 | 4.88 | <0.001 | -0.21 ± 0.41 | -0.52 | 0.61 |
|  | Variance explained by a random factor, ID = 0.09 |  |  | Variance explained by a random factor, ID = 0.08 |  |  |
| Saturation change |  |  |  |  |  |  |
| Age | 1.35 ± 1.35 | 1.00 | 0.33 | 5.21 ± 2.11 | 2.48 | 0.021 |
| Treatment | 7.65 ± 1.09 | 7.00 | <0.0001 | 2.68 ± 1.38 | 1.95 | 0.07 |
| Interaction | 6.56 ± 1.55 | 4.24 | <0.001 | -0.77 ± 1.95 | -0.40 | 0.70 |
|  | Variance explained by a random factor, ID = 2.86 |  |  | Variance explained by a random factor, ID = 11.40 |  |  |
| Brightness change |  |  |  |  |  |  |
| Age | -1.70 ± 2.06 | -0.83 | 0.42 | 5.08 ± 3.43 | 1.48 | 0.15 |
| Treatment | 11.48 ± 1.58 | 7.29 | <0.0001 | 2.77 ± 2.00 | 1.38 | 0.19 |
| Interaction | 13.74 ± 2.23 | 6.17 | <0.0001 | 0.86 ± 2.83 | 0.31 | 0.76 |
|  | Variance explained by a random factor, ID = 7.99 |  |  | Variance explained by a random factor, ID = 34.77 |  |  |

Bird identity (ID) was included as a random factor in each model.
Note that sample sizes were reduced in this analysis because we lacked measurements in two samples (one in the adult UV treatment group and the other in the young control group). For the paired experimental design (see Methods), we had to exclude “unpaired” samples as well.

**Figure S1.**
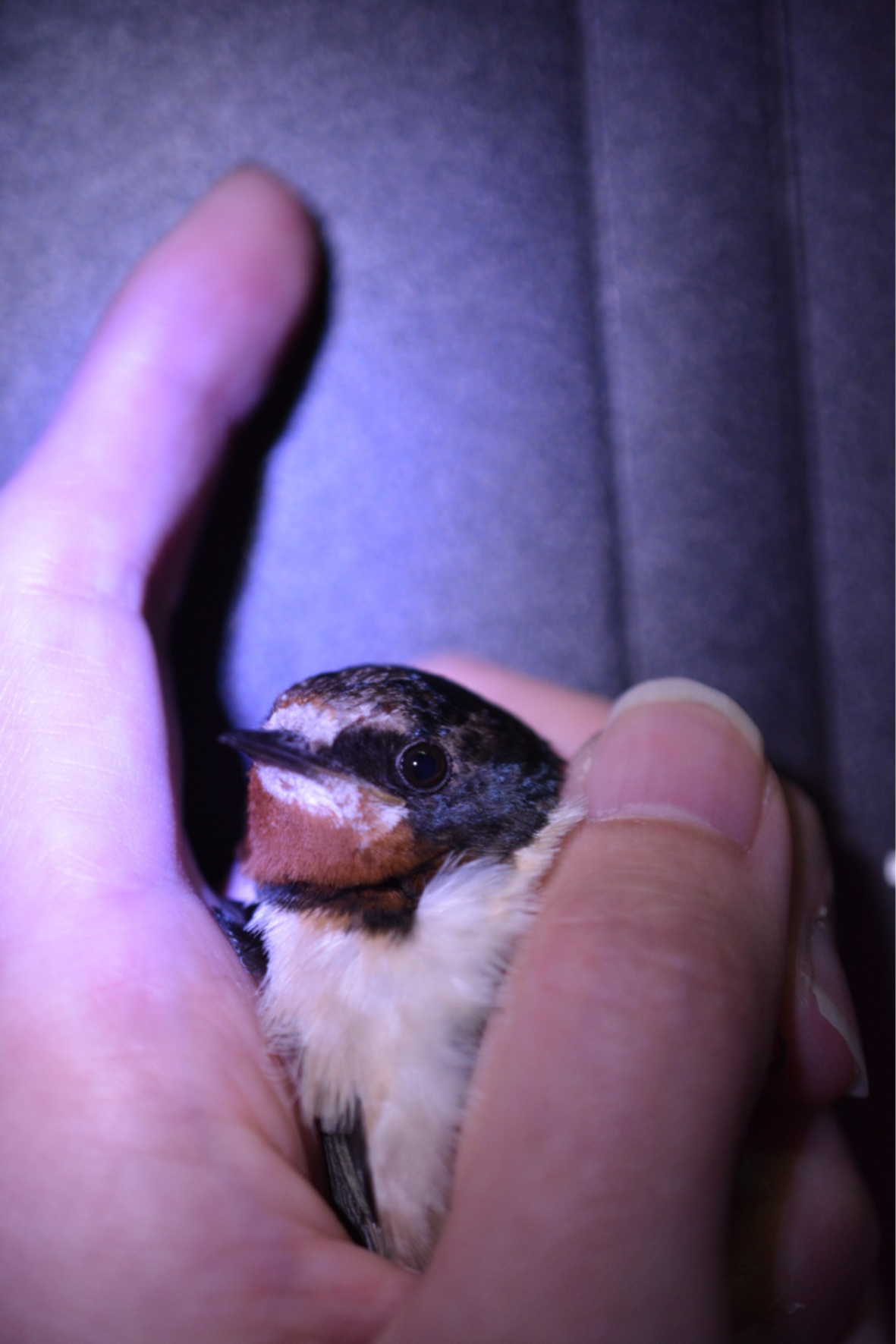
Reddish throat patch of a molting barn swallow. Note the color difference between newly molted (darker) and previously molted, faded (paler) feathers

**Figure S2.**
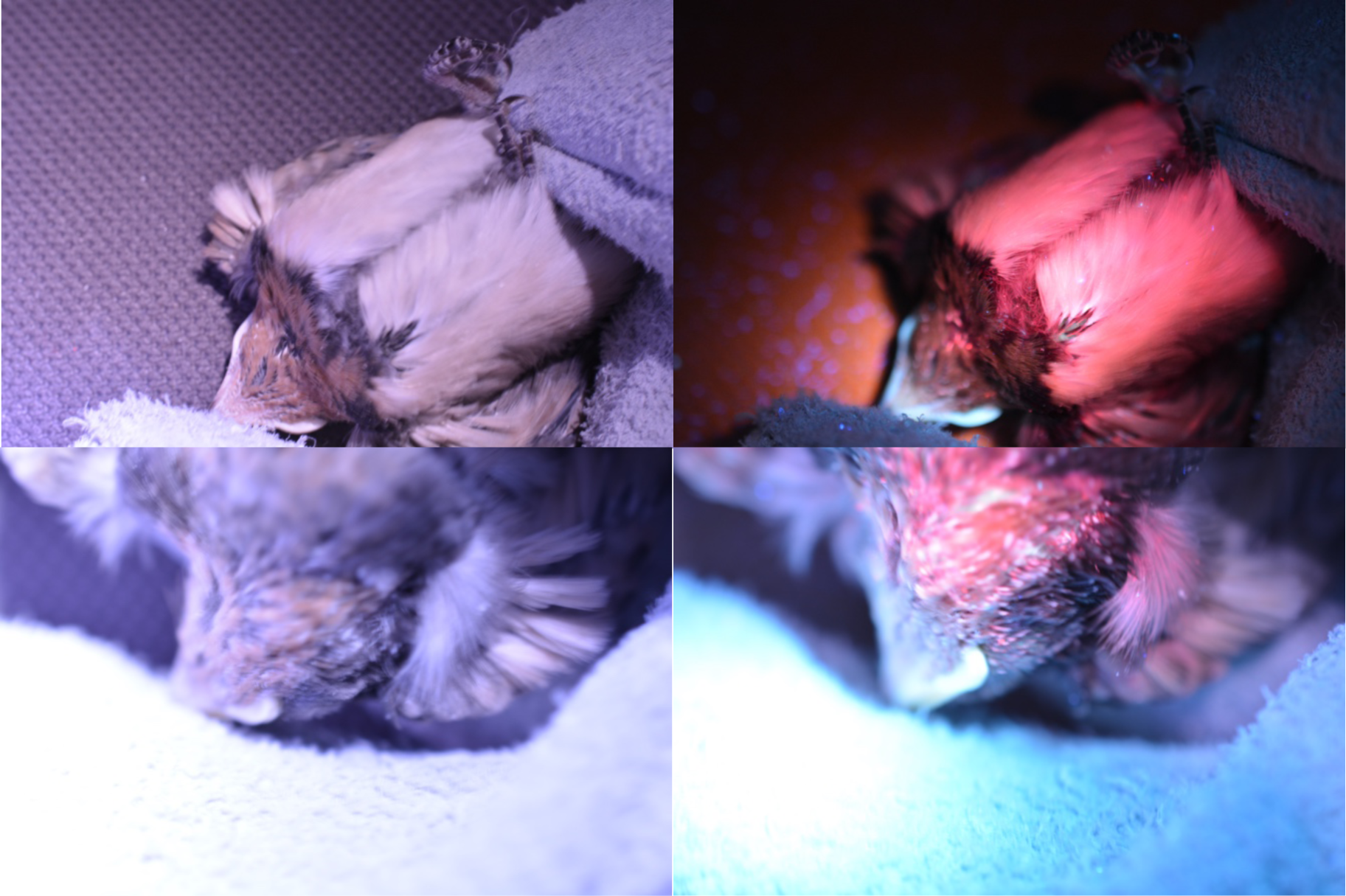
Photographs of a 15-day-old nestling barn swallow taken under white light and UV light (upper left; ventral surface under white light; upper right: ventral surface under UV light; lower left; throat region under white light; and lower right: throat region under UV light). Note the fluorescent red coloration under UV light

**Figure S3.**
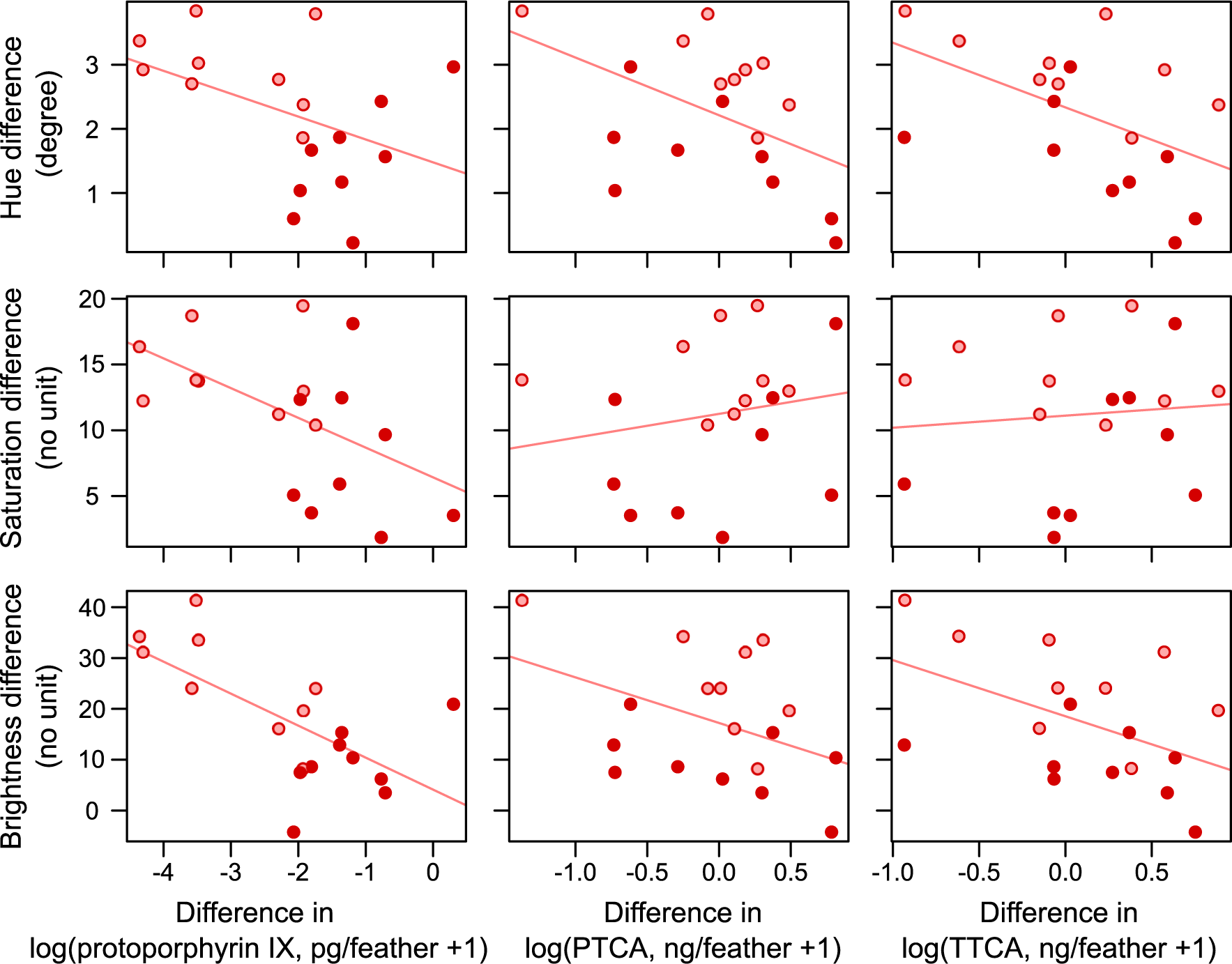
Relationships between color and pigmentation differences between treatments in the barn swallow: protoporphyrin IX (left column); PTCA (middle column); TTCA (right column). PTCA and TTCA are indices of eumelanin and pheomelanin, respectively. The upper, middle, and bottom rows show hue change, saturation change, and brightness change, respectively. Red and pink circles (dark and light gray, in print) indicate adults and young, respectively. Simple regression lines are depicted (see Table S4 for detailed statistics)

